# Representational geometry reveals how neuronal diversity supports perceptual performance

**DOI:** 10.1101/2025.06.26.661754

**Authors:** Sonica Saraf, J. Anthony Movshon, SueYeon Chung

## Abstract

A complete understanding of population coding requires connecting multiple levels of neural processing: individual responses, population representations, and behavior. We link these by relating the distribution of neuronal tuning properties to a population’s representational geometry and its efficiency for perceptual tasks. We use theory, analysis of recordings from macaque primary visual cortex (V1), and simulations to reveal how diversity of tuning amplitude and bandwidth enhances the population code for visual discrimination and identification. Both types of diversity drive different, but complementary changes to the representational geometry. Amplitude diversity increases the Euclidean distance between the responses to different stimuli, while bandwidth diversity creates a larger angular distance between them. The first utilizes the range of firing rates available to neurons, and the second exploits the high-dimensional nature of population responses. Population codes can be improved using these two different geometric changes, and amplitude and bandwidth diversity provide biological mechanisms for doing so.

**Highlights:**

- Perceptual performance is improved both by increased diversity of response amplitude and increased diversity of tuning bandwidth.
- Both kinds of diversity improve visual discrimination and identification.
- Amplitude diversity improves discrimination more, and bandwidth diversity improves identification more.
- Representational geometry reveals the mechanisms of these effects.

## Introduction

What makes neuronal representations efficient? Answering this question requires a framework that connects the properties of individual neurons to population-level representations and to behavioral performance. Such a framework should allow us to understand how neuronal response properties can be adjusted to maximize performance. Neurons exhibit varying levels of stimulus selectivity. Some are broadly responsive, while others are more specifically tuned (Eliav et al., 2021; Fitzgerald et al., 2006; Goris et al., 2015; Ringach et al., 2002). The functional role of this diversity remains unclear. Previous work by Shamir and Sompolinsky (2006) and Ecker et al. (2011) showed that tuning diversity affects coding for fine discrimination tasks. But how this diversity produces its population-level effect has remained unknown. To address this gap, we turn to representational geometry (Chung and Abbott, 2021), a recently introduced framework that captures how neural populations encode stimuli for downstream tasks. Here we ask what population-level geometric changes underlie the ways that diversity of neuronal tuning improves representational efficiency.

A neural population’s ability to code stimuli for downstream tasks is related to the structure of its responses, captured by its representational geometry. These population responses occupy a high-dimensional space in which each axis corresponds to one neuron’s firing rate. A point in this space represents the population’s response to a presentation of a stimulus. Points from various trials or slight variations in the stimulus (depending on the scientific question at hand) are grouped together and form a representation encoding the stimulus (DiCarlo and Cox, 2007; Roweis and Saul, 2000; Tenenbaum et al., 2000; Yerxa et al., 2023). Analyzing the geometry of these representations has emerged as a powerful technique for understanding population coding in many brain areas (Bernardi et al., 2020; Chung and Abbott, 2021; DiCarlo and Cox, 2007; Genkin et al., 2023; Mehrpour et al., 2021; Murray et al., 2017; Tian et al., 2024). More recently, an approach known as *manifold capacity* extended the classical notion of perceptron capacity (Cover, 1965) from single points to structured, manifold-shaped representations linking the task-relevant geometry of population responses to downstream decoding performance (Chung et al., 2018). The decoding assumption for capacity is separability – downstream populations measure the linear separability between the representations for various stimuli.

Here, we revisit the question of why tuning diversity is beneficial to population coding by measuring the capacity of both recorded macaque V1 populations and simulated populations. Capacity, measured in manifolds per neuron, reflects how efficiently a population uses its available neural dimensions to support separable representations: higher capacity means more stimuli can be distinguished by a downstream linear decoder using the same number of neurons, or equivalently, that the same number of stimuli can be distinguished using fewer neurons. Therefore, capacity measures the *representational efficiency* of a population for perceptual tasks. We create a new framework to relate the geometry of the full structure of representations, which is distinct from prior geometric characterizations of the worst-case points for linear separability, to their representational efficiency. We then use this framework to understand how tuning diversity alters geometry to enhance representational efficiency for perceptual tasks.

Specifically, we consider diversity in two response properties, amplitude and bandwidth, and its impact on representational efficiency for two different perceptual tasks, discrimination and identification. Fine discrimination is the detection of small stimulus differences, such as distinguishing a vertical and off-vertical orientation. Information coding for fine discrimination has been studied both theoretically and experimentally (Green et al., 1966; Jazayeri and Movshon, 2006; Seung and Sompolinsky, 1993) because it helps us understand the limits of our perception. We also consider identification – when one stimulus must be distinguished from all others. Identification has a special role in everyday vision, because it is an obligate part of programming a movement or executing a search: for example, reaching for the correct utensil, searching for a face in a crowd, reading. We focus on discriminating and identifying orientations so that we can study well-parameterized tuning curves. We take stimulus orientation as an early analog of objects and natural scenes: if tuning diversity impacts V1 population codes for orientation, the same principles should apply to coding for naturalistic stimuli in higher-level visual areas (Wang and Ponce, 2022).

We find that tuning properties map onto specific geometric properties: amplitude diversity expands the Euclidean distance between manifold centers (centroid norms), while band-width diversity reduces the angular similarity between them (center correlations). Through distinct geometric changes, each type of diversity enhances representational efficiency, but in ways that differentially benefit discrimination and identification. Our results suggest that population codes benefit from exploiting both geometric transformations, and that tuning diversity is a biological mechanism for achieving this. A population could achieve these same benefits by increasing the number of neurons, or amplitudes, etc., but tuning diversity does so while respecting metabolic constraints.

By linking neuronal tuning to representational geometry and efficiency, we show how tuning diversity alters population codes to aid perception. Tuning diversity may not be just a consequence of biological variability, but rather a fundamental population coding strategy for behavior. We offer a conceptual framework for viewing neuronal selectivity through the lens of representational geometry.

## Results

### Geometric framework

We examined how well populations of real and simulated V1 neurons distinguish the orientation of drifting sinusoidal gratings. To study how tuning diversity shapes representational efficiency, we analyzed population geometry, defining population-level representations in the same experimental paradigm we used to analyze tuning curves (Figure 1). The left side of Figure 1A shows the experimental design. Briefly, monkeys were shown 50 trials each of 72 drifting gratings that differed in their orientation (0 deg, 5 deg, …, 175 deg), for a total of 36 different orientations. Recordings were made with multi-electrode arrays placed in V1 of 5 anesthetized macaque monkeys (Graf et al., 2011). Our simulated population responses followed a similar paradigm (see *Methods*). The right side of Figure 1A shows schematics of neuronal tuning curves of various amplitudes and bandwidths, cartooning the diversity observed in V1 (Ringach et al., 2002).

**Figure 1.**
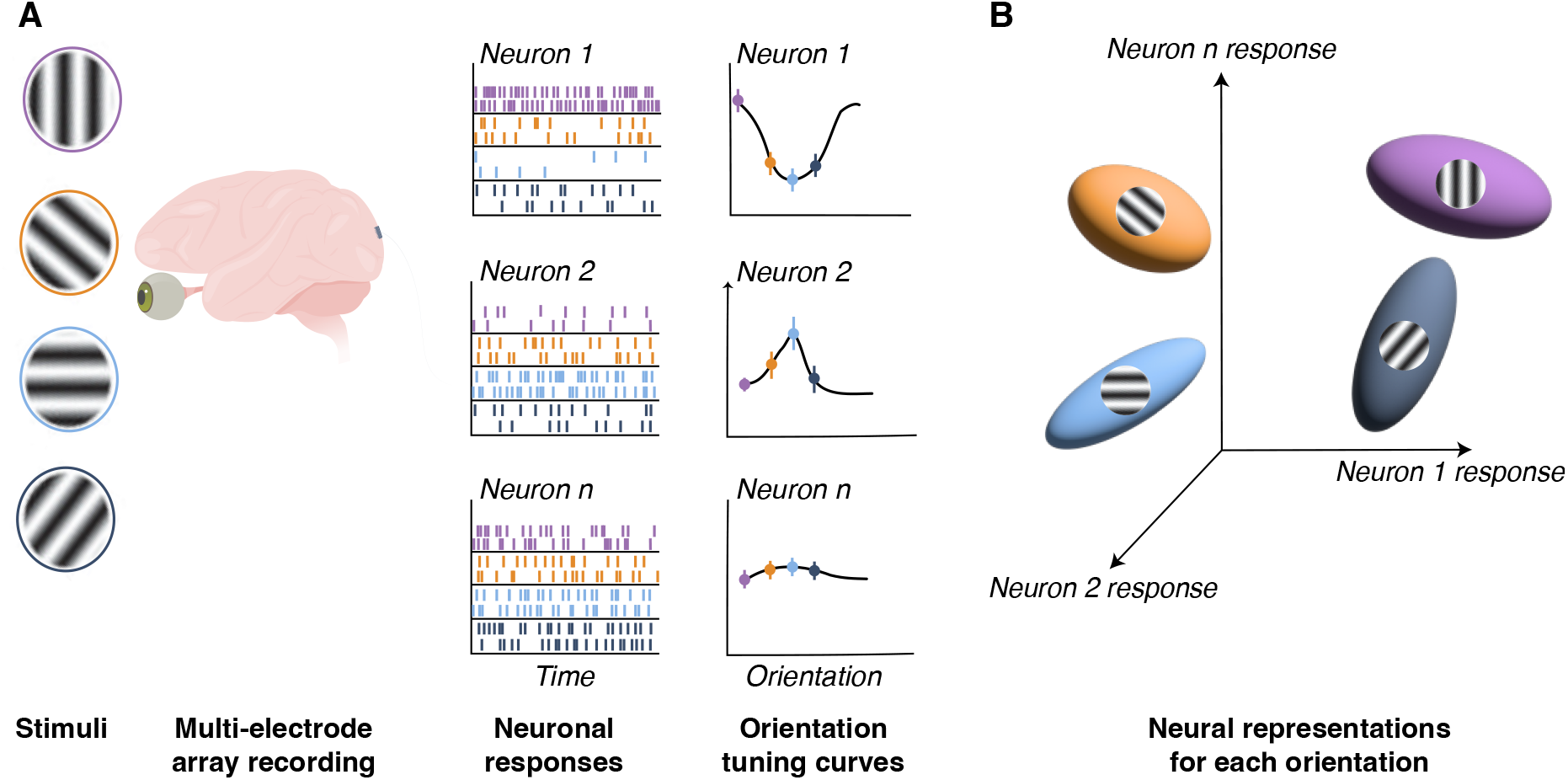
Schematic of tuning curves and neural manifolds. **A** Experimental procedure, simulated rasters and tuning curves. Neuronal data were previously reported in Graf et al. (2011), and we made analogous measurements in our simulations. A neuron’s average response to each condition gives a point on the neuron’s tuning curve. **B** The population’s responses to each stimulus condition form the neuronal manifold encoding that stimulus. A point on the manifold represents the population’s response to one trial, and the centers of the manifolds are the average responses to each stimulus (i.e. the values on the tuning curves).

We define the representations from the population’s trial by trial responses to each stimulus (Figure 1B). For a given orientation, we collect the population activity across every trial in which that orientation was presented; these points form a neuronal representation, or *neuronal manifold*) (Chung and Abbott, 2021). Presenting *P* different orientations for *S* trials each therefore yields *P* different manifolds, each containing *S* points. The shapes of the manifolds are determined by trial-to-trial variability and covariability in the population because each manifold encodes a single orientation. The population’s representational efficiency depends on the geometry of these manifolds and is captured by a single quantity, its *capacity*. Capacity reflects how many neuronal dimensions are needed to separate the *P* manifolds with hyperplanes – each from one nearby manifold (discrimination; Figure 2A) or from all of the other manifolds at once (identification; Figure 2B). Specifically, the capacity is the ratio of the number of manifolds to the minimum number of neuronal dimensions required to reach a threshold level of separability (Chou et al., 2025; Chung et al., 2018); see *Methods* for details.

**Figure 2.**
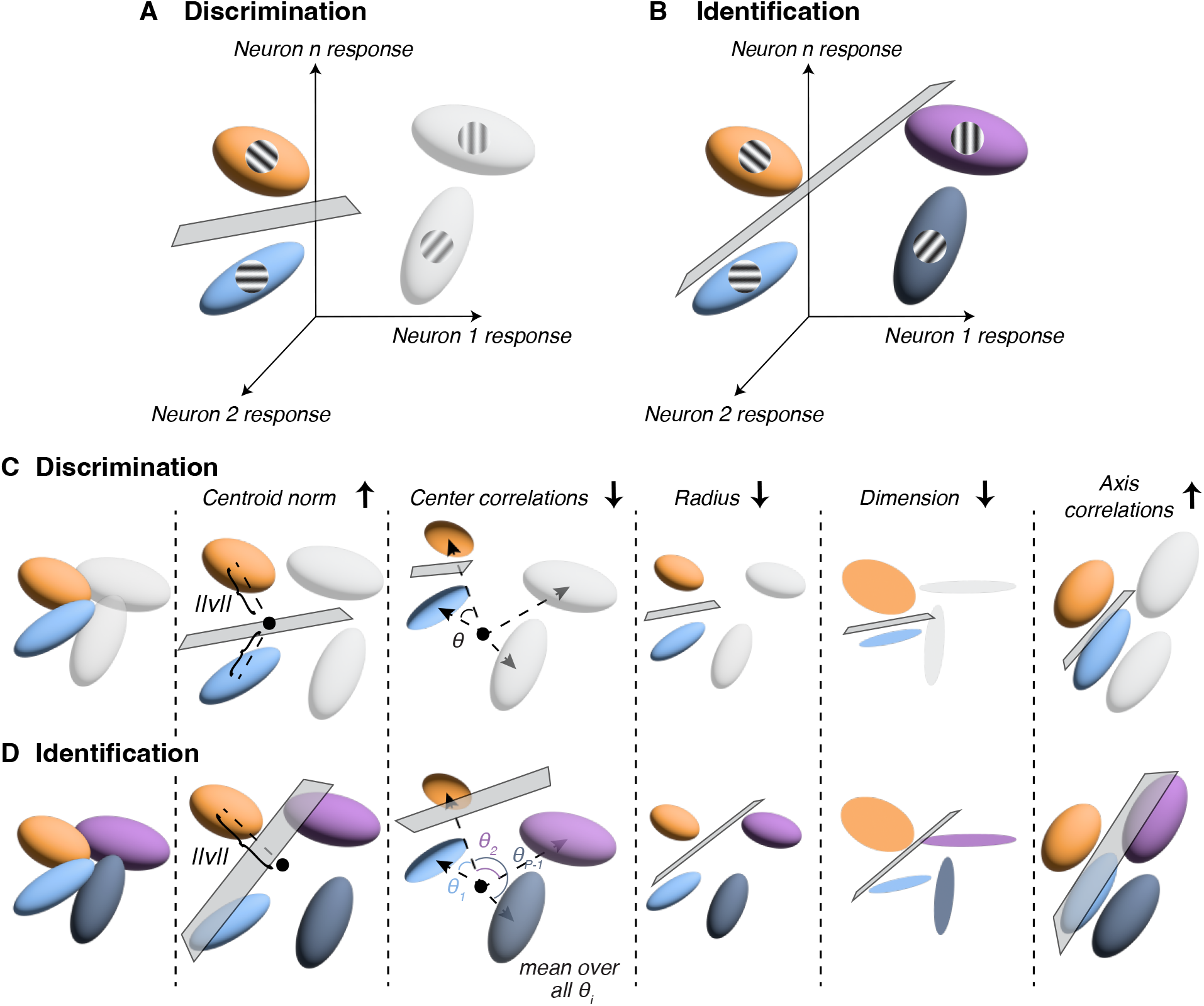
Tasks and geometric changes that increase capacity. **A** Discrimination task. Capacity, a measure of representational efficiency, is measured by linearly separating the manifolds for neighboring stimuli. **B** Identification task. Here, capacity is measured by linearly separating each manifold from the other *P* − 1 manifolds. **C** Geometric properties and their relations to capacity for both tasks. The first two properties measure aspects of the *signal* in the representations. The centroid norm is the average Euclidean distance (||*v*||), between the center of each of the two manifolds and the global mean of all of the manifolds, which is indicated by the black dot. Center correlations measure the angular distance between the centers of the two manifolds. The following three measures capture aspects of the noise in the representations. The radius and dimension measure the overall size and shape of the two manifolds being separated. The radius measures the average distance between a manifold’s boundary points and its center. The dimension measures the average projection of random vectors onto the manifold’s boundary points. Finally, the axis correlations capture the similarity of the principal axes of the two manifolds. **D** Analogous to panel C, but for the identification task. Here, the centroid norm, radius, and dimension are defined as they are for the discrimination task, except instead of averaging over the measures for the two manifolds to discriminate, we report the values for only the target manifold. The black dot indicates the global mean of all manifolds. Center correlations are the average of the angular distances between the target manifold’s center and every other manifold’s center. The axis correlations are an average of the angular distances between the target manifold’s axes and all other manifold’s axes. Throughout the paper we report the averages of the geometric measures over all possible fine discrimination and identification tasks. This equates the centroid norm, radius, and dimension across the two tasks.

#### Geometric properties and their relations to capacity

##### Geometric properties

We developed a set of geometric properties designed to bridge neuronal response properties and population-level representations. Previous work using capacity, such as Chung et al. (2018) and Chou et al. (2025), measures the geometry of the worst-case points for linear separability. Our geometric properties instead characterize the full point-cloud structure of manifolds, which makes the connection between response properties and geometry more intuitive.

Consider first the discrimination task. Figure 2C shows five geometric properties and how each shapes representational efficiency for discriminating the orange manifold from the blue. The first subpanel shows shows a starting configuration, and the following subpanels reflect geometric changes that improve representational efficiency. The first two properties, the centroid norm and center correlations, capture the signal separation between the representations. The centroid norm is the Euclidean distance between a manifold’s mean point (its “center”) and the global mean of the responses to all *P* stimuli (black dot); manifolds farther from the global mean have a larger centroid norm. The center correlations measure the angle between the manifolds’ center vector directions; manifolds that occupy more distinct subspaces of the response space have lower center correlations. The remaining three properties characterize the noise in the representations. The radius and dimension are properties measuring the average size and shape of each individual manifold, which arise from trial-to-trial response variation: a smaller radius reflects lower variability, and a flatter manifold has lower dimension. Finally, the axis correlations quantify the average angle between manifolds’ principal components; greater alignment of the manifolds’ orientations corresponds to higher axis correlations.

An analogous set of geometric measures exists for the identification task, which we implement as discriminating one target manifold from all the others. The two tasks differ in one key respect: identification geometry must account for the global structure of all manifolds. Figure 2D shows the five geometric properties for identifying the orange manifold (the target) from the rest. We define the centroid norm, radius, and dimension in the same way as for discrimination, except that instead of averaging the measures over the two manifolds being separated, we only consider the target manifold’s measures. Throughout the paper, we report the average value of the properties over identification for all possible targets (and all possible pairs for fine discrimination). This makes the centroid norm, radius, and dimension the same across the two tasks. On the other hand, for identification the center and axis correlations are defined to measure global structure: in the third and last subpanels of Figure 2D, the center and axis correlations are averages of the pairwise center and axis correlations, respectively, between the target and *all* other manifolds.

##### The relationship between geometric properties and representational efficiency

Previous work has linked these geometric properties and capacity, but those links rest on the worst-case point geometry (Chou et al., 2025; Chung et al., 2018; Wakhloo et al., 2023). We instead established analogous relationships between our geometric measures and capacity empirically, as follows:

Using a set of simulated data with a large range of neuronal tuning parameters, we performed a multiple regression among the five geometric properties and capacity to infer the geometric changes that improve representational efficiency. The coefficients for the multiple regression are shown in the first two columns of Table 1. These coefficients establish the relationship between geometry and representational efficiency, which are illustrated in panels C and D of Figure 2.

**Table 1.** The first two columns show the multiple regression results establishing the relationships between geometric properties and capacity for discrimination and identification. We used 50 simulations each of 25 different diversity types – all combinations of amplitude and bandwidth diversity levels: 0, 2.2, 5, 11.8, and 23 spikes/s for amplitude diversity and 0, 2.8, 5.7, 11.0, and 15.5 deg for bandwidth diversity. We computed the multiple regression over all 1,250 simulations and show the standardized coefficients for each of the geometric properties as predictors of capacity. The next four columns show the predicted capacity increase (percentage) due to the changes in each geometric property from the homogeneous to most diverse population in our simulations. The middle two columns show the results for the amplitude diversity manipulation and the final two show the results for the bandwidth diversity manipulation. Notice that the centroid norm predicts the largest increase in capacity for amplitude diversity, and the center correlations predict the largest increase for bandwidth diversity.

| Geometric property | Regression $\beta$ | | Amplitude | | Bandwidth | |
| --- | --- | --- | --- | --- | --- | --- |
|  | Disc. | Ident. | Disc. | Ident. | Disc. | Ident. |
| Centroid norm | 0.87 | 0.62 | 83.0 | 79.0 | -21.5 | -20.5 |
| Center correlations | -0.88 | -0.75 | -10.1 | -33.6 | 89.1 | 78.0 |
| Radius | -0.11 | -0.37 | -1.49 | -7.03 | 5.43 | 25.7 |
| Dimension | -0.066 | -0.038 | 4.91 | 3.74 | -1.42 | -1.08 |
| Axis correlations | -0.011 | 0.26 | -0.55 | -2.63 | -0.011 | 22.1 |

A higher centroid norm improves representational efficiency because it will increase the Euclidean distance between manifolds. Lower center correlations also improve representational efficiency by increasing the angular distance between manifolds. Smaller manifolds are less likely to overlap, so they are easier to separate from each other. Thus, lower radii and dimensions increase representational efficiency. Finally, higher axis correlations generally increase representational efficiency.

### Observations from neuronal data: amplitude diversity

Previous work has analyzed how amplitude diversity influences Fisher information in populations with noise correlations. Using simulated data and theoretical analysis, these studies found that amplitude diversity can overcome information limitations imposed by to noise correlations (Ecker et al., 2011; Shamir and Sompolinsky, 2006). Motivated by this finding, we wanted to check if amplitude diversity also improves population coding in our macaque V1 data.

We analyzed previously collected microelectrode array recordings from V1 of four hemispheres from three macaques (Graf et al., 2011). To confirm the analytical results from previous work, we first measured the relationship between amplitude diversity and Fisher information. From each recorded population, we drew subpopulations with varying levels of amplitude diversity: exploring a range of sample sizes, we found that 500 subsets of 30 neurons each provided sufficient diversity while keeping the subpopulations large enough (the data set in Figure 3 contained 59 cells). We excluded occasional outlying cells to keep the statistics reasonable.

**Figure 3.**
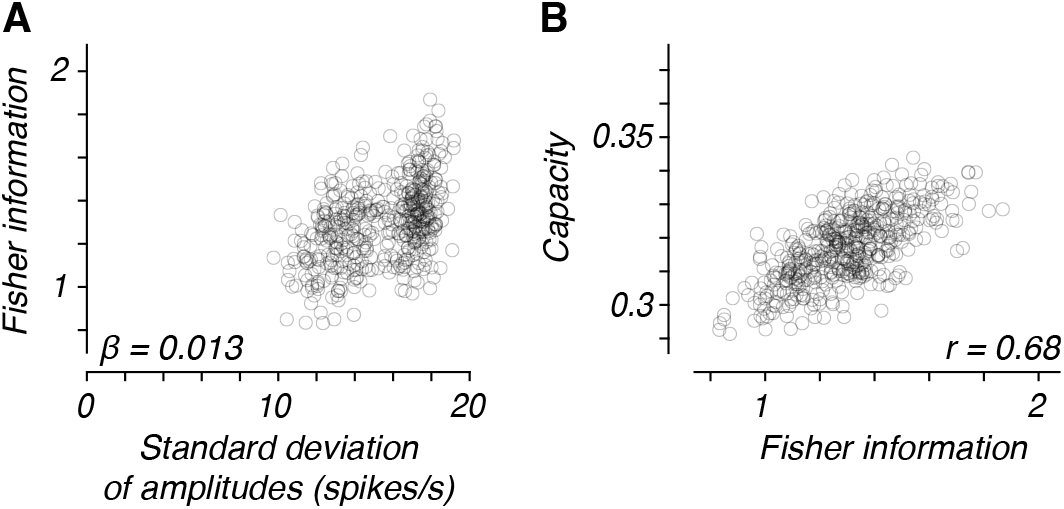
The relationships between amplitude diversity, Fisher information, and capacity for fine discrimination in neuronal populations from macaque V1. We randomly chose 500 subpopulations of 30 neurons each from the full population. **A** Fisher information and versus the amplitude diversity for the subpopulations. The inset shows the multiple regression coefficient between the amplitude diversity and Fisher information in the subpopulations. **B** Capacity for discrimination versus the Fisher information for the same subpopulations as in panels A. The inset shows Pearson’s correlation between capacity and Fisher information for the subpopulations.

Figure 3A shows the relationship between amplitude diversity and the Fisher information of these subsets. To parse how much of any increase in information came from amplitude diversity, we performed a multiple regression between the standard deviation of the amplitudes, the mean amplitude, and the Fisher information of each subset. The standardized regression coefficient for the amplitude standard deviation explaining Fisher information was *β* = 0.013, indicating that amplitude diversity had a negligible effect on information. This could be due to other uncontrolled variations across subsets, such as different noise correlations and tiling of the stimulus space, which are controlled in simulations like those in Shamir and Sompolinsky (2006) and Ecker et al. (2011). For these same subsets, we measured representational efficiency using capacity, the measure that we will focus on for the rest of the paper. Capacity for discrimination and Fisher information were correlated (Figure 3B, Pearson’s correlation coefficient of 0.68), validating that capacity is a reasonable measure of discriminability.

Next, we studied the relationship between amplitude diversity and capacity. Recall that amplitude diversity did not predict the increase in Fisher information, likely due to uncontrolled variations across subsets, such as uneven tiling of the stimulus space and different noise correlations. To address these issues, we created pseudo-populations that evenly spanned the full range of orientation preferences (see *Methods*). This procedure corrected the uneven tiling evident in the first subpanel of Figure 4C, and by reassigning trial-by-trial responses across stimuli, it also removed noise correlations. Figures 4A and B show the relationship between amplitude diversity and capacity for true and pseudo-populations, respectively. We again performed a multiple regression between the standard deviation of the amplitudes, the mean amplitude, and the capacity for discrimination. We display the multiple regression coefficients in the first two columns of Table 2. Amplitude diversity had a strong impact on capacity for discrimination for both true subpopulations (*β* = 0.47) and pseudo-populations (*β* = 0.66).

**Table 2.**
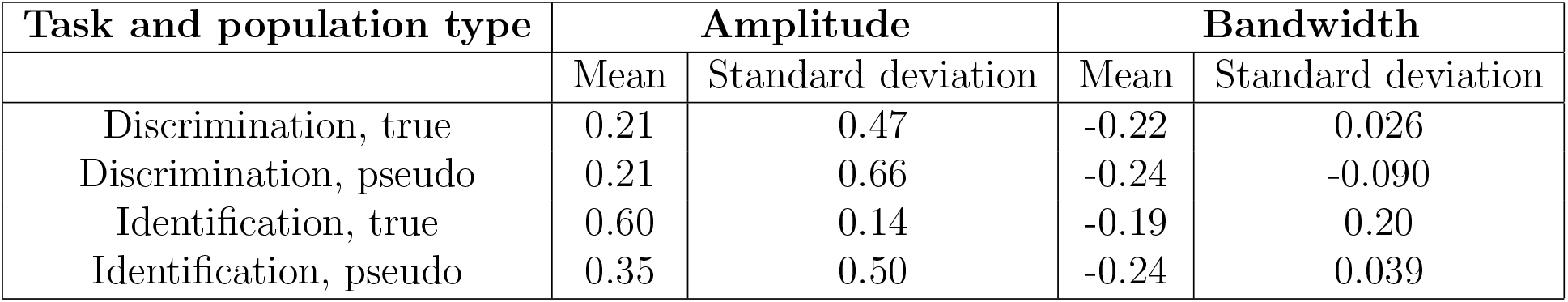
Multiple regression results for subpopulation properties as predictors of capacity. From data set 3, we created 500 subpopulations and 500 pseudo-populations. We measured the mean amplitude, standard deviation of amplitudes, mean bandwidth, and standard deviation of bandwidths for each of the subpopulations, and their capacity for discrimination and identification. For each task and type of subpopulation, we show the standardized regression coefficients for the mean and standard deviation of the tuning properties as a predictors of capacity.

**Figure 4.**
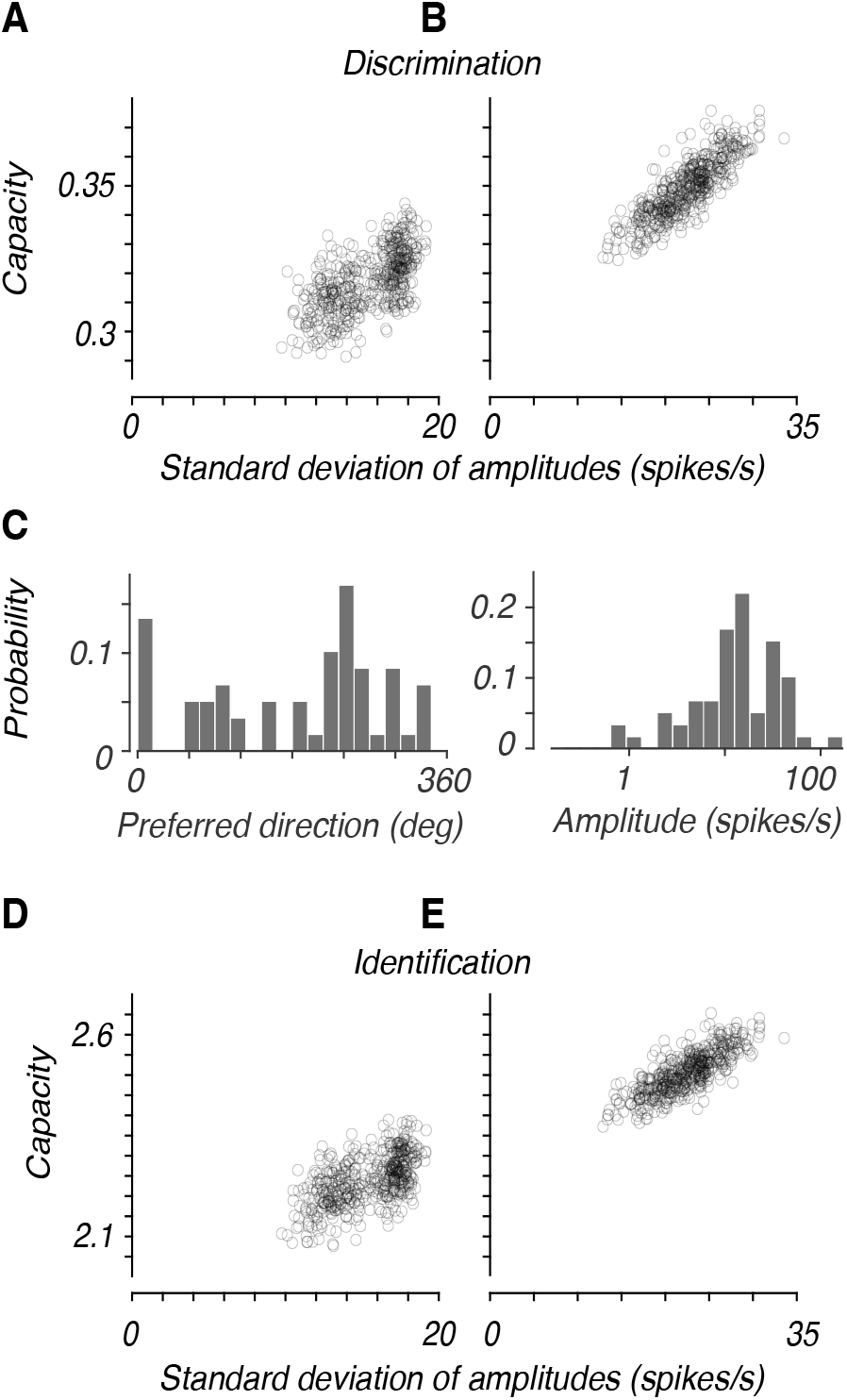
The relationships between amplitude diversity and capacity in neural populations from macaque V1. In panels A and D, 500 subpopulations of 30 neurons each were chosen from the full population, excluding an outlier. Without excluding the outlier, the subsets would have clustered into a group of low amplitude diversity and a group high diversity. In panels B and E, we used 500 upsampled, pseudo-populations of 180 neurons each. **A,B** Capacity for fine discrimination versus the amplitude diversity for the subpopulations and pseudo-populations, respectively. **C** Distribution of preferred stimuli and peak to trough amplitudes of the cells in the data set (data set 3 from Graf et al. (2011)). See the Methods section for the definition of peak to trough amplitude. Observe from the left panel that the neuronal preferences do not evenly tile the stimulus space. We created the upsampled pseudo-populations in panels B and E for even tiling of orientation preferences. We did not exclude the outlier cell with high amplitude for the pseudo-populations of B and E since it was likely that each of the 59 cells were selected at least once per population. **D,E** Capacity for identification versus the amplitude diversity for the subpopulations and pseudo-populations, respectively. Note that amplitude diversity has a stronger effect for discrimination capacity than for identification capacity.

Fisher information is defined only for discrimination, but capacity also measures representational efficiency for identification. Amplitude diversity increased capacity for identification in both populations as well (Figure 4D, *β* = 0.14 for true populations, and 4E, *β* = 0.50 for pseudo-populations), though less strongly than it increased discrimination capacity.

Together, these analyses indicate that amplitude diversity improves representational efficiency for discrimination and identification. A parallel analysis of bandwidth diversity was inconclusive (regression coefficients in the last two columns of Table 2; relationship between capacity and diversity for all four data sets in SI Figure 2, with multiple regression coefficients in SI Tables 1-4). Bandwidth diversity ranged from a standard deviation 5.0 to 15.5 degrees for the full data sets, but see the range of values occupied along the abscissa for each individual cloud of points in SI Figure 2C and D. This indicates that the subpopulation diversity fell in a much smaller range.

The neuronal data have several limitations, including small population sizes, uneven tiling of the stimulus space, and limited ranges of bandwidth diversity across subpopulations. To control these variables precisely, we turn next to simulated populations.

### Amplitude and bandwidth diversity for discrimination and identification in simulations

We modeled neuronal tuning curves as von Mises functions. The preferred stimuli of the neurons were evenly spaced across orientations (180 deg). In the homogeneous populations, the tuning curves for all cells had the same amplitude. Empirical observations show lognormally distributed amplitudes in macaque V1 (see the distribution of amplitudes in Figure 4C and SI Figure 1B). So, to create populations with diverse amplitudes, we randomly chose each neuron’s tuning curve amplitude from a lognormal distribution whose mean matched the homogeneous population. To create higher levels of diversity, we increased the standard deviation of the lognormal distribution. We simulated five levels of diversity with standard deviations of the lognormal being 0 (homogeneous), 2.2, 5, 11.8, and 23 spikes/s. We selected this range to span the variability in our four neuronal datasets, whose amplitude standard deviations were 10.5, 12.4, 23.0, and 8.6 spikes/s.

We introduced bandwidth diversity into neural populations in a similar fashion, but used gamma distributed bandwidths – again, in imitation of experimental observations (see SI Figure 1C). The five levels of bandwidth diversity corresponded to standard deviations of 0, 2.8, 5.7, 11.0, and 15.5 deg. As with the range of standard deviations for amplitude diversity, we selected an experimentally reasonable range for bandwidth standard deviations (the standard deviations of the bandwidths for the four arrays were 5.0, 8.3, 9.2, and 15.5 deg).

We simulated the responses of populations of 300 neurons to 50 trials each of 36 different oriented drifting gratings (5 degree spacing to match the sampling in the neuronal data). We assumed a Poisson-like noise model for each cell– meaning that the response variance was the same as the mean response rate, and a population noise correlation that decayed with the difference in the preferred directions for a pair of cells (Shamir and Sompolinsky, 2006).

For each population, we measured the geometry and representational efficiency for discrimination and identification. We analyzed: 1) the relationship between diversity and representational efficiency for each perceptual task, 2) the relationship between diversity and the geometric properties for each task, and 3) the geometric properties that explained the changes in representational efficiency. In Figure 5 we show the relationship between each of the five geometric properties and capacity for the different levels of amplitude and bandwidth diversity.

**Figure 5.**
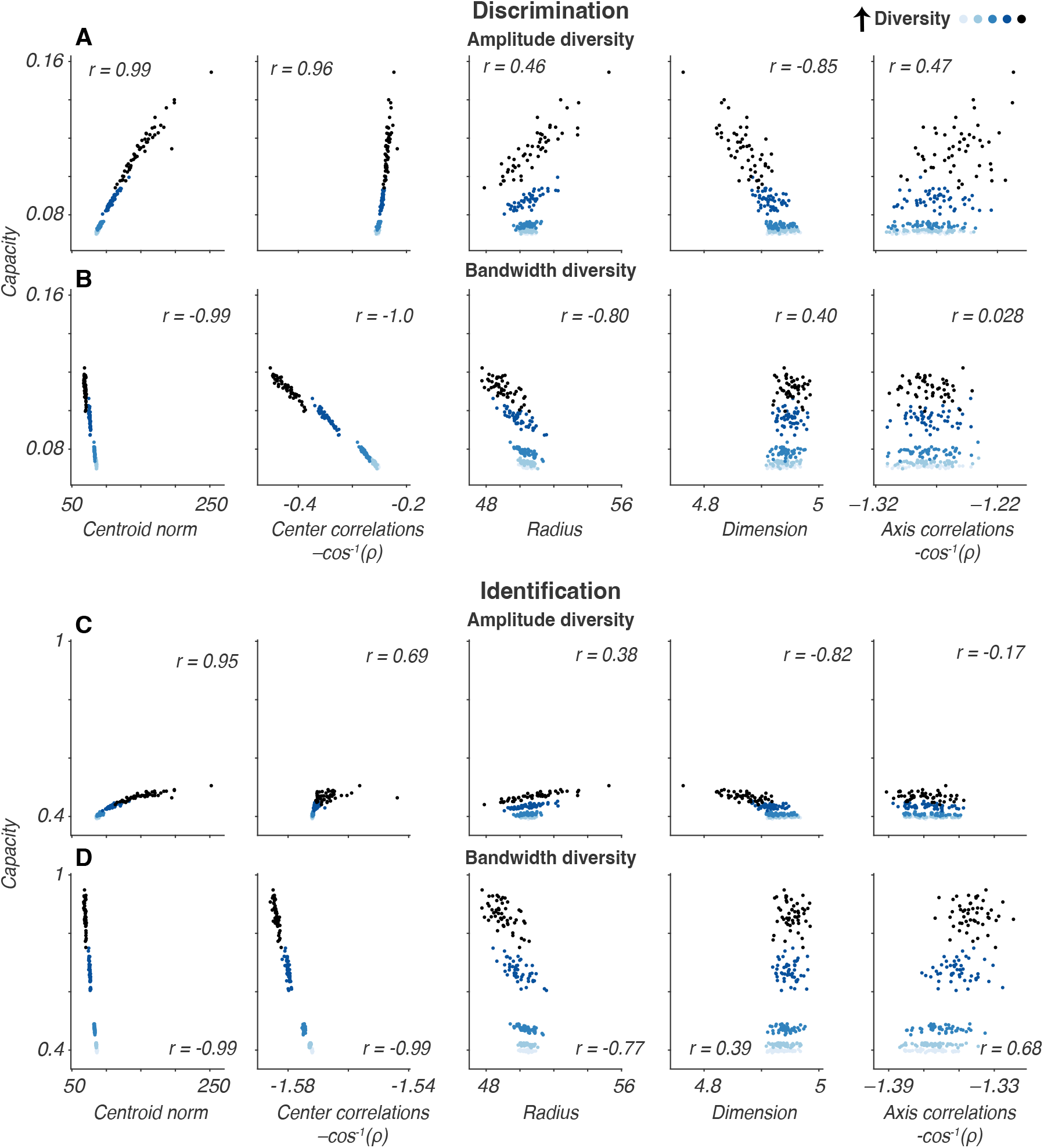
Scatter plots showing the relationships between geometric properties and capacity as tuning diversity changes in simulations. From lightest to black, the colors indicate increasing levels of diversity (standard deviations of amplitudes of 0, 2.2, 5, 11.8, and 23 spikes/s and standard deviations of bandwidths of 0, 2.8, 5.7, 11.0, and 15.5 deg). Each panel lists the Pearson’s correlation coefficient between the geometric property and capacity. **A** Geometry versus capacity for amplitude diversity simulations and discrimination. **B** The same as panel A, but for bandwidth diversity and discrimination. **C** Amplitude diversity and identification. **D** Bandwidth diversity and identification.

#### Diversity improves representational efficiency

Consider the first subpanel of Figure 5A to understanding the results from our simulations. For each of the 5 diversity levels indicated in the legend, we plot the centroid norm versus capacity for discrimination for 50 populations. First, note the relationship between diversity and capacity. The homogeneous populations (lightest colored dots) have lower capacity than the most diverse population (black dots). This trend can be seen for amplitude diversity and discrimination (Figure 5A), bandwidth diversity and discrimination (Figure 5B), amplitude diversity and identification (Figure 5C), and bandwidth diversity and identification (Figure 5D). So, across both types of diversity and tasks, capacity increases from the homogeneous to most diverse case.

For discrimination, homogeneous populations had an average capacity of 0.071 manifolds per neuron. This means that ≈ 28 neurons from a homogeneous population are needed to perform discrimination. Comparing the average of the capacity values for the lightest colored and black dots in Figure 5A, we find that there was a 61% increase in capacity for discrimination due to amplitude diversity. This increase corresponds to 0.0434 additional separable manifolds per neuron, which means that the most diverse population can encode for discrimination with 10.7 fewer neurons than the homogeneous population.

In Figure 5C, we show the relationship between capacity and geometry for identification as amplitude diversity changes. Homogeneous populations had an average capacity of 0.40 manifolds per neuron, which means that ≈ 90 neurons from a homogeneous population are needed to perform identification. Note that in general, capacity for identification is higher than for discrimination because there are 36 manifolds being separated rather than 2, and some of these are much easier to separate from the target manifold. Capacity for identification increased with amplitude diversity by 18%. This corresponds to 0.0712 manifolds/neuron and 13.7 fewer neurons needed to encode for identification with the most diverse population.

We next turn to bandwidth diversity (Figures 5B, discrimination; 5D, identification). The ordinates are matched with panels A and C, respectively, to aid comparing the effects of amplitude and bandwidth diversity. There was a 57% increase in capacity for discrimination due to bandwidth diversity (0.04 additional separable manifolds/neuron – 10.2 fewer neurons needed for the most diverse population; see Figure 5B). This is smaller than the 61% increase due to amplitude diversity, and one can see this by noting the higher range of values along the ordinate for the plots in Figure 5A in comparison to Figure 5B.

For identification, bandwidth diversity raised capacity by 117% (0.465 manifolds/neuron, 48.8 fewer neurons needed for the most diverse population; see Figure 5D). This gain exceeds the 18% from amplitude diversity, as the lower range in Figure 5C versus Figure 5D shows.

#### Diversity’s impact on geometry

We measured the changes in each of the five geometric properties attributable to changes in diversity. These are indicated by the change along the abscissa across the differently colored dots in each scatter plot of Figure 5. The abscissae for each geometric property are the same for Figures 5A and B (discrimination) and for Figures 5C and D (identification). This allows us to compare the relative change in each property due to the different diversity types.

The first two properties, centroid norm and center correlations, show the cleanest relationships with diversity level. For example, in the first panel of Figures 5A and C, the differently colored dots, which represent different diversity levels, occupy different ranges of the abscissa. This indicates that amplitude diversity impacts the centroid norm, and specifically, observe that amplitude diversity increases the centroid norm. See *Supporting Information* for a formal proof that the centroid norm increases with amplitude diversity. In the second panel of Figures 5B and D, we see an analogous segregation of center correlation values due to bandwidth diversity. In this case, bandwidth diversity decreases center correlations. The other three properties do not show as much consistency in their relationships with diversity.

#### Geometric changes underlying improvements in representational efficiency

Having identified which geometric properties diversity affects, we asked whether the changes in these geometric properties explain the gain in representational efficiency from the homogeneous to the most diverse populations. To address this quantitatively, we calculated the percentage change in capacity that was predicted by the change in each of the five geometric properties. To do so, we found the changes in the properties from the homogeneous to most diverse case and multiplied them by the beta coefficients established from the regression results shown in the first two columns of Table 1. We used the unstandardized coefficients to calculate predicted increases in capacity. The predicted percentage increases in capacity are shown in the last four columns of Table 1. For both discrimination and identification, the centroid norm predicts the largest capacity increase from amplitude diversity (see the third and fourth columns of Table 1): amplitude diversity improves efficiency by moving the manifold centers farther apart. Center correlations predict the largest capacity increase from bandwidth diversity (see the last two columns of Table 1), indicating that bandwidth diversity improves efficiency by decorrelating manifold centers.

### Comparing amplitude and bandwidth diversity

Our simulations reveal that amplitude and bandwidth diversity improve representational efficiency through distinct transformations to representational geometry. Figure 6 illustrates these geometric transformations. The center column shows a homogeneous population’s tuning curves and geometry. The left side shows how amplitude diversity affects the tuning curves and geometry. Amplitude diversity pushes representations for different stimuli further apart by increasing their centroid norms. The right side shows how bandwidth diversity affects the tuning curves and geometry. Bandwidth diversity increases the angular separation between manifolds. Intuitively, one can interpret the larger centroid norms as utilizing the full range of firing rates available to the population, while the larger angular separation utilizes the high dimensional nature of the population.

**Figure 6.**
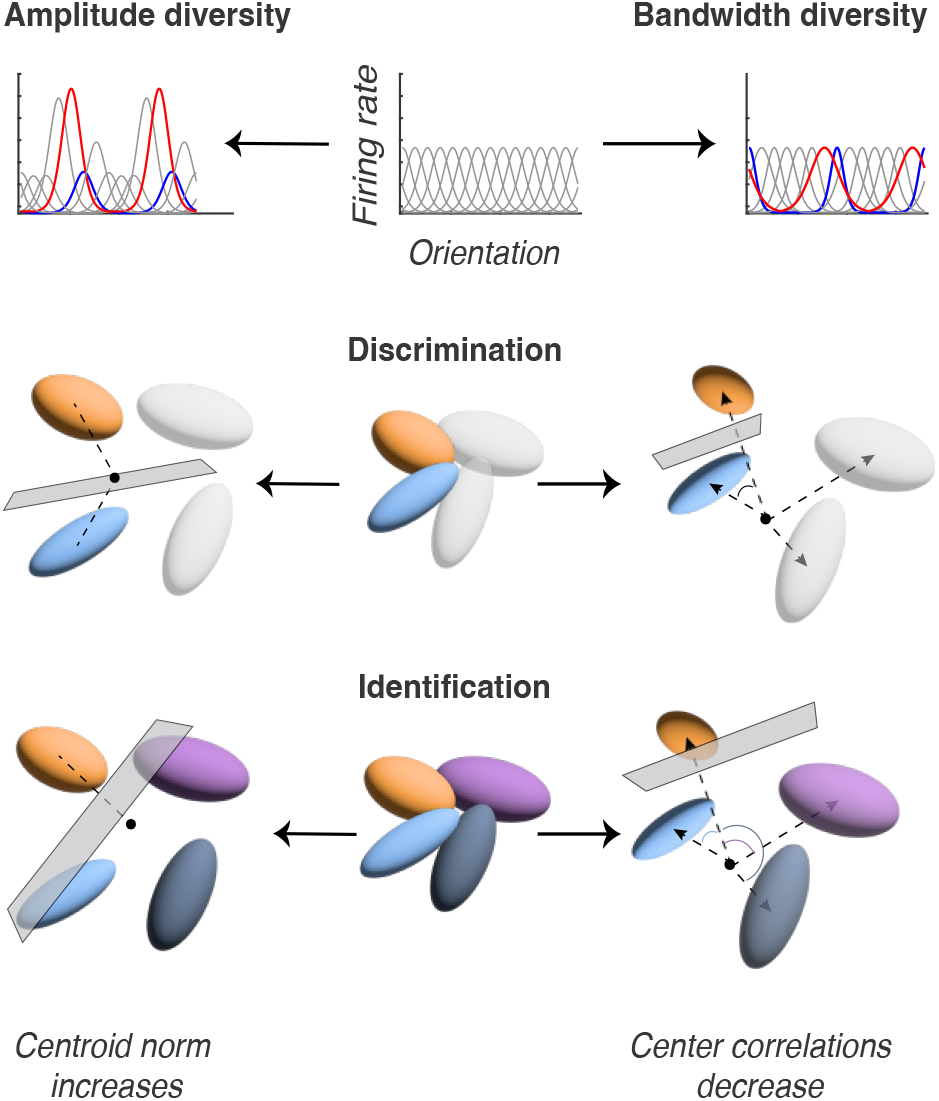
Schematic illustrating the geometric transformations caused by either type of diversity. Amplitude diversity increases the Euclidean distance between manifolds while bandwidth diversity lowers the center correlation by expanding the angular distance between manifolds.

Both types of diversity increase representational efficiency for both the discrimination and identification tasks. However, each diversity type favors one task: impacts one of the two tasks more: amplitude diversity improves discrimination more, while bandwidth diversity improves identification more (compare the lines where one diversity type is homogeneous between Figures 7A and B). To understand why this occurs, we look to the percentage changes in capacity predicted by each geometric property in the last four columns of Table 1. The main improvements to capacity are due to the centroid norm (under amplitude diversity) and center correlations (under bandwidth diversity). In addition, across the two tasks, their effects are roughly the same. So, what causes the differential impacts on the two tasks? It must be due to changes in the remaining geometric properties. As we will see in what follows, identification is more sensitive to the *global* structure of the manifolds than discrimination, and bandwidth diversity gives a larger benefit to global structure than amplitude diversity does.

**Figure 7.**
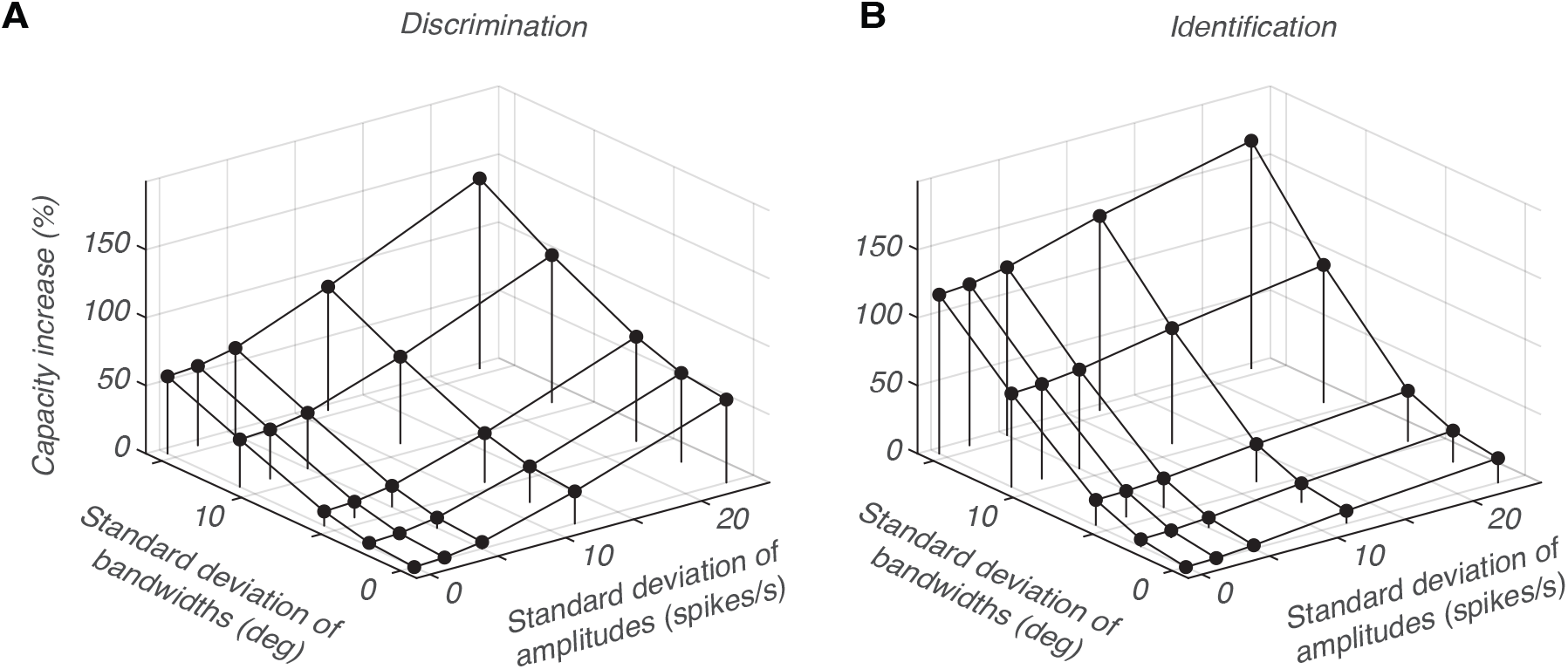
How tuning diversity affects capacity for discrimination and identification. We take the average capacity over all 50 simulation trials at each of the 25 combined diversity levels to calculate the changes in capacity shown. The 25 diversity levels are all of the combinations from the standard deviation of amplitudes of 0, 2.2, 5, 11.8, and 23 spikes/s and the standard deviation of bandwidths of 0, 2.8, 5.7, 11.0, and 15.5 deg **A** Change in capacity for discrimination. Filled circles show the average percentage increase in capacity from the homogeneous case for the 25 diversity combinations for the discrimination task. Open circles show the same but for the identification task. **B** Same as panel A, but for identification. Note that both types of diversity increase capacity for discrimination by roughly the same amount, but bandwidth diversity improves capacity for identification far more than amplitude diversity does. Additionally, the two types of diversity have approximately independent effects on capacity.

Why does amplitude diversity improves discrimination more? The coefficients table (Table 1) shows that the change in center correlations under amplitude diversity predicts a larger capacity decrease for identification than for discrimination. Recall that the center correlations are defined differently for the two tasks: we consider only the closest pairs when defining the correlations for discrimination, but for identification, we consider the correlations between all other manifolds and the target. Amplitude diversity causes a larger increase in the center correlations among all the manifolds than for just nearby pairs (0.019 radian increase for discrimination versus 0.18 radians summed across all 35 pairs for identification), which explains why the change in the center correlations predicts a sharper decrease in capacity for identification.

For an analogous reason, the change in axis correlations due to bandwidth diversity predicts a larger increase for identification. Bandwidth diversity increases the axis correlations among all the manifolds more than for just the nearby pairs (≈ 0 radians increase for discrimination versus 0.83 radians across all 35 pairs for identification). This is part of the reason why bandwidth diversity helps identification more than discrimination. Comparing the final two columns in Table 1, we also see that the change in radius predicts a larger increase for identification than for discrimination. The radius is defined exactly the same for the two tasks, so it changes by the same amount for both tasks. This suggests that identification benefits more than discrimination does from an overall decrease in the size of the manifolds. Because identification relies on separating one manifold from many, it depends on noise in many more directions than discrimination does, which only depends on noise in the direction between two manifolds. Thus, a decrease in radius, which indicates a noise reduction across all directions, will improve representational efficiency for identification more.

So far, we have considered amplitude and bandwidth diversity separately. However, biological neural populations exhibit both types of diversity, which raises the question of how amplitude and bandwidth diversity interact. In Figure 7, we show the capacity increase from the homogeneous population to populations created with 25 different combinations of amplitude and bandwidth diversity: all combinations created from the set of standard deviation of amplitudes of 0, 2.2, 5, 11.8, and 23 spikes/s and the set of standard deviation of bandwidths of 0, 2.8, 5.7, 11.0, and 15.5 deg. We compared the capacity of these populations with the capacities of the populations with only amplitude or bandwidth diversity. We found an approximately independent relationship between capacity improvements due to amplitude and bandwidth diversity for both discrimination and identification. This confirms that amplitude and bandwidth diversity create fundamentally different geometric changes. Altogether, this result suggests that populations can combine both types of diversity to maximize their representational efficiency.

### Diversity improves representational efficiency under nuisance variation

Natural stimuli vary in many ways that are irrelevant to the task at hand, requiring an observer to ignore nuisance variables while performing specific perceptual tasks (e.g. determining the orientation while ignoring variations in size or shape). Adding such a variable, such as contrast, enlarges the manifolds encoding orientation, because the same orientation now elicits a wider range of responses. We expect that tuning diversity would allow these larger manifolds to have higher representational efficiency than they would if the population had homogeneous tuning.

We tested this idea by introducing contrast dependent responses and randomized stimulus contrasts into our simulations, and measuring how diversity affected the capacity for discrimination and identification. For both discrimination and identification, homogeneous populations had lower capacity when we added random contrast fluctuations. When we introduced amplitude and bandwidth diversity, the capacity matched and exceeded the capacity for the simulations without nuisance variables (compare the open circles above amplitude and bandwidth diversity with the filled circle above homogeneous for both panels in Figure 8). Interestingly, however, amplitude diversity did not improve representational efficiency for discrimination under contrast variations as much as it did without– 61% increase without contrast nuisance (compare the first two filled circles in Figure 8A) and 20% with (compare the first two open circles in Figure 8A). All other improvements across diversity and task types were of roughly the same magnitude as in the no nuisance variation case.

**Figure 8.**
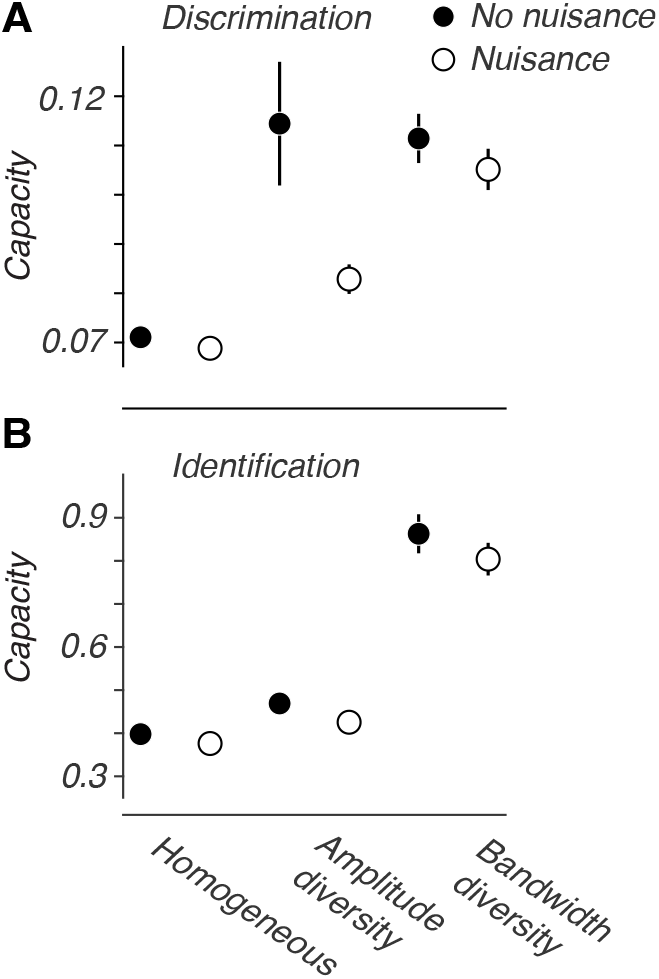
Diversity improves representational efficiency in the presence of nuisance variation. **A** Capacity for discriminating orientation in the presence of contrast variations. Introducing contrast variations lowers capacity, but capacity is restored by tuning diversity in the population. We plotted points indicating the homogeneous and most diverse cases from the simulations in Figure 5 for comparison (filled circles). Error bars indicate the standard deviation over 50 simulations. Note that amplitude diversity doesn’t improve capacity as much as it did in the nonuisance case. The amplitude diverse population and bandwidth diverse population had respective standard deviations of 23 spikes/s and 15.5 deg. **B** Same as panel A, but for identification. The range of capacity values spans the same ratio as that in panel A.

Why did amplitude diversity not improve representational efficiency for discrimination under contrast variations as much as it did without? Contrast variations interact with the neuron’s amplitude of response because they scale the firing rates of neurons. One can show that these variations effectively lower the overall amplitude diversity of the population when measured from the responses across the stimuli: if *A*_*i*_ is the *i*th neuron’s amplitude, with an average contrast of *c*, the measured amplitude will be *Ã*_*i*_ = *cA*_*i*_. The variance of *Ã*_*i*_ is given by 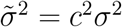, where *σ*^2^ is the variance of *A*_*i*_. If we take *c* = 0.5, which it approximately was for these simulations, 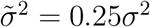.

This reduced diversity explains why amplitude diversity falls short of the benefit it would provide without contrast variation. We expect that bandwidth diversity would be similarly impaired when faced with nuisance variations that scale tuning widths, such as when manifolds encode stimuli with dispersions about a main orientation. Even so, both types of diversity improved representational efficiency under contrast variations for both discrimination and identification. This suggests that tuning diversity may therefore be a mechanism for flexibly encoding a specific stimulus variable in the presence of others that are irrelevant to the current task.

## Discussion

What properties of the neuronal response can improve representational efficiency? We address this question by connecting neuronal tuning properties, population-level representational geometry, and decoding for perceptual tasks. Our central result is that diversity of neuronal tuning improves population coding through two complementary geometric transformations: it increases the distance between representations and reduce their angular similarity.

Diversity of selectivity is common across neural systems, including hippocampal place field sizes in bats (Eliav et al., 2021) and rats (Rich et al., 2014), finger pad selectivity in macaque somatosensory cortex (Fitzgerald et al., 2006), head-direction tuning bandwidths in mouse postsubiculum (Clark et al., 2025; Duszkiewicz et al., 2024), retinal selectivity in salamander (Berry II et al., 2019), and frequency tuning in rat auditory cortex (Insanally et al., 2019). Our results reveal that this widespread diversity is not merely biological variability, and it can systematically reshape population representations and enhance representational efficiency. Moreover, amplitude and bandwidth diversity improve coding via distinct mechanisms: amplitude diversity increases the distance between stimulus representations in the neural activity space, while bandwidth diversity increases their angular separation by decorrelating the directions of their centers. Both transformations (by amplitude and bandwidth diversity) increase separability, but they do so in ways that differentially affect different perceptual tasks. Amplitude diversity has the stronger effect on fine discrimination, whereas bandwidth diversity has the stronger effect on identification.

To illustrate the complementary nature of the geometric transformations, imagine the region of population activity space occupied by the stimulus representations as a box enclosing the representations: a larger box can hold more separable representations, and its size can increase in two ways, by lengthening its sides or by gaining additional independent dimensions (as in the difference between a flat envelope and a three-dimensional box). By expanding the centroid norms of the manifolds, amplitude diversity lengthens the box, allowing the population code to use more of the firing-rate range available to its neurons. Bandwidth diversity, on the other hand, spreads the manifold centers across more independent directions, effectively increasing the dimensionality of the box and allowing the code to exploit the high dimensionality of population response space. Amplitude and bandwidth diversity therefore expand the space containing the representations geometrically distinct ways. Together, they allow more stimulus representations to remain separable than either form of diversity would allow alone.

These geometric benefits could also be achieved by increasing the number of neurons or their firing rates, but either change would increase metabolic cost. Tuning diversity offers a different solution: it improves the representational efficiency while keeping total spiking approximately constant. On average, the total number of spikes emitted by the simulated population remained nearly unchanged as amplitude diversity increased (4.13, 4.12, 4.12, 4.12, and 4.13×10^6^ spikes from the homogeneous to most diverse populations). As bandwidth diversity increased, total spiking fell slightly (4.13, 4.13, 4.12, 4.06, 3.92×10^6^ spikes from the homogeneous to most diverse populations). Thus, neural systems can enhance representational efficiency without increasing metabolic cost by distributing spikes unevenly across neurons. This parallels efficient-coding accounts, in which limited channel capacity drives heterogeneity in how a population allocates its coding resources (Jun et al., 2022). Indeed, under reasonable assumptions, a diverse population will *always* have at least as much discriminability as a homogeneous population with the same number of spikes (Ringach, 2026). This raises a further question: how do neuronal circuits generate and maintain tuning diversity? Future work could examine how diversity changes over evolution, development, and learning.

Previous work has established that tuning diversity can benefit neural coding. For example, Shamir and Sompolinsky (2006) and Ecker et al. (2011) showed that tuning heterogeneity can overcome information saturation in populations with noise correlations (Panzeri et al., 2015), and that the resulting information gain depends on the strength of those correlations. These studies focused mainly on how amplitude diversity affects fine discrimination. More recently, Tian et al. (2024) showed that firing-rate diversity improves fine discrimination by lowering the dimensionality of population covariability. Together, these studies show that diversity can improve discrimination, but they leave open two questions central to the present work: whether the benefits of diversity extend to identification, and whether different forms of diversity improve coding through distinct geometric mechanisms. Our representational geometry approach addresses both questions. It allows us to compare discrimination and identification within a common framework and to show how amplitude and bandwidth diversity *differentially* shape representational efficiency in the population code.

In our framework, the manifold size is determined by trial-to-trial variability and noise correlation (Pillow et al., 2008), raising the question of whether our results depend on the Poisson-like noise model in the main simulations. They do not. When we repeated the simulations with additive noise, we observed the same main effects: amplitude diversity increased centroid norms and helped discrimination more, while bandwidth diversity decorrelated manifold centers and helped identification more. Under Poisson-like noise, diversity also affected radius, dimension, and axis correlations, which characterize noise in the neuronal responses. These effects arise because Poisson-like noise scales with firing rate. Under additive noise, where variability does not scale with firing rate, these noise-related geometric properties did not change.

We also tested whether the benefits of diversity persist when the population must encode one stimulus variable while ignoring variation in another. To do so, we introduced contrast variations into the stimulus manifolds and found that both amplitude and bandwidth diversity still improved capacity. This result moves the simulations toward more naturalistic settings, where behavior often depends on one stimulus parameter even as other, task-irrelevant parameters vary. Such nuisance variation makes the manifolds more complex than simple point clouds, which poses a challenge for classical approaches to population coding, including linear Fisher information, that depend on assuming specific assumptions about response distributions. The manifold geometry and capacity framework used here avoids those assumptions. By characterizing the full geometric structure of the representations, it allows us to quantify representational efficiency even when nuisance variation distorts the shape of these population response.

Recent work has used geometry to study population coding with multiple stimulus variables (Wakhloo et al., 2026), and has shown that diversity of selectivity to task-relevant variables can improve linear readout (Posani et al., 2025). Our work addresses a complementary question: how does tuning for a single variable shape representational geometry. We focused on V1 populations coding stimulus orientation because orientation tuning in V1 is well-characterized, allowing us to connect neuronal tuning properties directly to population-level geometry and representational efficiency. In doing so, our work bridges the gap between analytical approaches grounded in single neuron tuning and approaches based on population-level geometry (Kriegeskorte and Wei, 2021).

Geometric analysis also reveals why the benefits of tuning diversity depend on the task. Identification depends more strongly than discrimination on the global structure of signal and noise because each target manifold must be separated from many alternatives, not just from a nearby manifold. As a result, changes that affect population responses across many directions have a larger impact on identification. For example, reduction in manifold size (radius) improves identification more than discrimination because identification is more sensitive to noise in all directions. Bandwidth diversity therefore provides a larger benefit for identification by improving the global structure of the manifolds, while amplitude diversity provides a larger benefit for discrimination by improving local separability between nearby manifolds. More broadly, different behavioral demands may favor different representational geometries, and different neuronal response properties may provide biological routes for achieving them.

By linking tuning properties to specific geometric transformations, our work suggests a general way to study neural population coding: identify the representational geometry required by a task, then ask which neuronal response properties produce that geometry. This perspective connects single-neuron tuning, population-level representational structure, and behavior within a common framework, and may help explain why diverse selectivity is so widespread across neural systems.

## Acknowledgements

We thank Albert Wakhloo, Nga Yu Lo, Artem Kirsanov, Will Slatton, Jenelle Feather, and Dario Ringach for helpful discussion about the work. We also thank Chi-Ning Chou for discussion and a well-managed codebase for measuring capacity (Chou et al., 2025). The Flatiron Institute is a division of the Simons Foundation. The computations reported in this paper were performed in part using resources made available by the Flatiron Institute. This work was supported in part by grants from the National Institute of Health– 5T32EY007136, 5T90DA059110, and R01DA059220 – and the Simons Foundation, Collaboration on the Global Brain (543019). S.S. is partially supported by an NYU GSAS Dissertation Fellowship. S.C. is partially supported by a Sloan Research Fellowship and a Klingenstein-Simons Award.

## Methods

### Definitions of geometric measures

We define geometry that captures the point-cloud structure of the manifolds and their relation to one another in the neural firing rate space. The geometry is designed to capture various aspects of the manifolds that relate to representational efficiency while maintaining intuitive relationships with tuning distributions. Therefore, the geometry is not the same as those from previous work using capacity (Chou et al., 2025; Chung et al., 2018), which characterizes the worst-case points for linear separability. Below, we define the geometry for both the discrimination and identification tasks.

#### Discrimination

Suppose that there are *N* neurons in the population, let *p* = 1, 2 represent the two manifolds to separate, and suppose that all manifold points have been centered with respect to the overall mean of the *P* manifolds. *C*_*p*_ is the center point of the manifold *p*. For each manifold *p*, we create a set of random Gaussian vectors {*t*_*i,p*_: 1 ≤ *i* ≤ *T*} where *T >> N*. If *t*_*i,p*_ is the *i*th Gaussian random vector originating at *C*_*p*_, we defined *s*_*i,p*_ to be the point on manifold *p* where *t*_*i,p*_ has the largest projection. This can be thought of as the closest boundary point for the manifold in the direction of *t*_*i,p*_. The vectors *a*_*i,p*_ are the unit vector principal components of manifold *p* and the coefficient *E*_*i,p*_ is the proportion of variance explained by component *a*_*i,p*_. Geometric definitions: **centroid norm**, ||*C*_*p*_||;**center correlations**,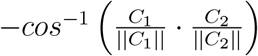 ; **radius**, 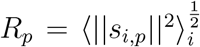 ; **dimension**, 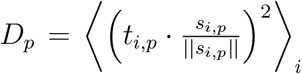 ; **axis correlations** 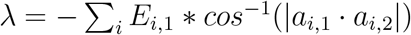

We negate the angle the correlations to preserve that lower center correlations improve separability – which was established in previous work on capacity theory (Wakhloo et al., 2023) that used different geometric definitions.

We averaged over ||*C*_*p*_|| , *R*_*p*_, and *D*_*p*_ to obtain the average centroid norm, radius, and dimension for the two manifolds in the discrimination task. Note that the trends we found with these radius and dimension definitions are similar to the trends shown with more conventional measures such as the *L*^2^ norm of the eigenvalues for radius and the participation ratio of the eigenvalues for dimension. However, we chose to use these definitions for radius and dimension to capture the sizes of non-ellipsoidal shapes.

For the values reported in Figure 5, we averaged the geometric measures over all possible fine discrimination tasks (between stimuli with 5 degree difference).

#### Identification

As before, suppose that there are *N* neurons in the population and that all manifold points have been centered with respect to the overall mean of the *P* manifolds. The definitions of *C*_*p*_, *t*_*i,p*_, *s*_*i,p*_, *a*_*i,p*_, and *E*_*i,p*_ are the same as for discrimination. Identification differs from discrimination because we are concerned with separating manifold *M*_*p*_ from the other *P* − 1 manifolds. This means that the global structure of all of the manifolds matters for identification, and which is reflected in the definitions for the center and axis correlations.

Geometric definitions: **centroid norm**, ||*C*_*p*_|| as above; **center correlations** are defined as: 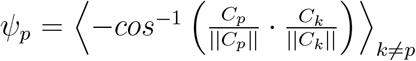 ; **radius**, *R*_*p*_ from above; **dimension**, *D*_*p*_ from above; **axis correlations**, 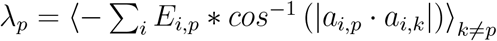

For discrimination, we measure only one pair of manifold’s correlations, while for identification we average over all possible pairs with the *p*th manifold.

For the values reported in Figure 5, we averaged geometric measures over all possible manifolds as *p*, the target manifold. Note that because we are averaging over all possible target manifolds (or nearby pairs for discrimination), the centroid norm, radius, and dimension end up being the same for both tasks.

### Measuring capacity

We measured capacity directly by following the methods described in Chung et al. (2018); Cohen et al. (2020) and improved upon in Chou et al. (2025). We briefly describe the procedure here: Suppose we have a population of *N* neurons and their responses to *P* different stimuli. For simplicity, we will describe the algorithm with reference to a specific discrimination task, separating the manifold for *θ*_1_ from the manifold for *θ*_2_. Capacity is empirically measured by finding the smallest *N*_*c*_ such that the manifolds for *θ*_1_ and *θ*_2_ are separable for 50% of the random projections of the data onto neural subspaces of dimension *N*_*c*_. We perform binary search to find *N*_*c*_, and capacity is measured to be 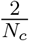, where is 2 is the number of manifolds to separate. The units for capacity are manifolds/neuron. We measure capacity analogously for identification for *θ*_1_, but instead measure the separability between the manifold for *θ*_1_ and ℳ:= ⋃_*i* ≠ 1_*Mi*. Capacity is 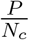 since there are *P* manifolds to separate.

Above we described measuring capacity for a specific discrimination or identification task. Since there are *P* = 36 possible discriminations (or identifications) in our simulations, we report the mean of the capacity values for the *P* discriminations of *θ*_*i*_ and *θ*_*i*+1_. For the edge case of *i* = *P* , we measure the separability between the manifolds for stimulus *P* and the first stimulus because orientation is a circular variable. We report an analogous averaged capacity for identification on our simulated data. Capacity is higher for identification than for discrimination because all *P* manifolds are being separated, and the further away manifolds are easier to separate from the target manifold.

#### Subsampling points for the neuronal data capacity measurements

To measure capacity, the manifolds must be linearly separable. Often times, when using all 50 trials per manifold in the neuronal data, the manifolds were not separable, likely because there were significantly fewer neurons per population than were in the simulated populations. To combat this issue, we downsampled to a random 8 of the 50 trials per manifold. We selected 8 trials as the largest number at which manifolds were separable across all data sets. We then averaged the capacity over 10 downsamplings. Note that the range of capacity values in Figure 5A for the simulated data is 0.07 to 0.11 which is lower than the 0.29 to 0.38 range for the neuronal data in Figure 4A,B. We confirmed that this is due to the manifolds having all 50 points instead of 8 as we used in the neuronal data analysis, because the capacity for simulations with 8 points per manifold is 0.28 to 0.34.

### Multiple regression analysis

We performed multiple regression analyses using the five geometric properties as explanatory variables and capacity as the dependent one. In the first two columns of Table 1 we report the standardized coefficients for all multiple regressions, 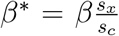, where *β* is the original coefficient and *s*_*x*_ and *s*_*y*_ are the standard deviations of the geometric property and capacity, respectively (Kerlinger and Pedhazur, 1973). We used the regular (unstandardized) *β*s when we calculated the predicted increase in capacity due to the changes in geometry caused by diversity.

### Simulated neural populations

#### Tuning curves

We followed the tuning curve model from Shamir and Sompolinsky (2006). The *i*th neuron’s tuning curve follows the function

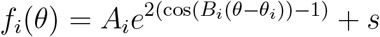

where *s* is the baseline firing rate for all neurons and *θ*_*i*_, *A*_*i*_, and *B*_*i*_ are the preferred stimulus, amplitude, and bandwidth parameters for neuron *i*, respectively. For a population of *N* neurons, the preferred stimuli were evenly distributed following the formula 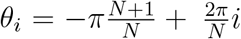. For all simulations shown, *s* = 3.3. The *f*_*i*_ are modified to create orientation selective cells such that *f*_*i*_ has two equally sized peaks 180 deg apart.

In the homogeneous populations, *A*_*i*_ = 15 and *B*_*i*_ is chosen such that the half-bandwidth of the cells is 16 deg. The half-bandwidth is defined as half of the width of the tuning curve at 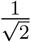 of the baseline-subtracted peak of the curve. For example, if the peak of the tuning curve is 15 + *s*, the half-bandwidth would be measured between the points where the tuning curve has the value 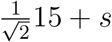.

We implemented amplitude diversity by assuming the amplitudes are lognormally distributed with a mean of 15 spikes/s. We create more diversity by increasing the variance of the lognormal distribution from which the *A*_*i*_ are selected. Please refer to the figures for the specific diversity levels used in the simulations.

Bandwidth diversity was introduced in a similar fashion, however we assume a gamma distribution for bandwidths. The mean of the half-bandwidths is set to 16 deg to match our biological neural populations, and bandwidths are drawn as follows:

Let *κ*_*b*_ be a scale parameter in {0.5 ,1, 2}, and select 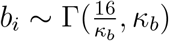. The expected value of *b*_*i*_ is 16, and the variance is 16*κ*_*b*_. Note that as *κ*_*b*_ increases, the population’s bandwidth heterogeneity increases. We then set

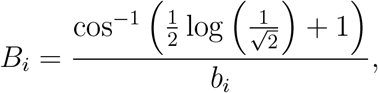

which gives *f*_*i*_ the desired half-bandwidth of *b*_*i*_. Throughout the text, we use the term “bandwidth” to mean the “half-bandwidth” as defined above.

#### Noise and correlations

Trial-to-trial variability and noise correlations are introduced in a similar manner to the method used in Shamir and Sompolinsky (2006). Correlations between neurons depended on the distance between their preferred stimuli, and exponentially decayed as the distance increased. However, we used multiplicative noise while Shamir and Sompolinsky (2006) assumed an additive noise model for their cells. The correlation between neurons *i* and *j* where *i* ≠ *j*, was given by 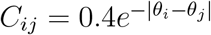.

We assumed Poisson-like statistics, and enforced that the trial-to-trial variability of each cell was equal to it’s mean response to a given stimulus. We then scaled the correlation matrix by the neural variabilities to create the covariance matrix, 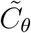, where *θ* indicates the stimulus direction.

Finally, we simulated the population’s responses on a trial of stimulus *θ* as 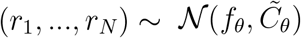, where *f*_*θ*_ = (*f*_1_(*θ*), …, *f*_*N*_ (*θ*)).

#### Contrast response functions

To model contrast dependent responses, we redefined *f*_*i*_(*θ*) as

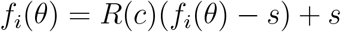

where 0 ≤ *c* ≤ 1 and 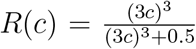. The parameter *c* scales the stimulus’s contrast, and we chose the function *R* to match the shapes of contrast response curves in V1 (Albrecht and Hamilton, 1982).

#### Contrast as a nuisance variable

For each trial response, we uniformly selected *c* from the range of 0.183 to 0.35 which ensured that *R*(*c*) fell within the range of 0.25 to 0.7, with a mean of 0.5. We also doubled the mean amplitude of our populations so that the mean responses were consistent with the simulations that did not have contrast variations.

### Neuronal data analyses

The data used here were previously reported in Graf et al. (2011). Neuronal data were recorded with Utah arrays implanted in V1 of 5 hemispheres of 3 macaques. Animals viewed drifting sinusoidal gratings of 72 different directions with 50 trials each. The spatial and temporal frequencies of the stimuli were chosen to maximize the population activity. The gratings were presented for 1,280 ms with a 1,280 ms blank screen in between stimuli. Data were spike sorted, and neural responses to the stimuli were taken from all 1,280 ms of each presentation. Further details of the data collection can be found in Graf et al. (2011). One array, data set 1, was reported by the authors of Graf et al. (2011) to have been under an altered experimental regime. Through our analyses, we also noticed a lower number of cells and lower overall firing rates in this data set than the other data sets showed. For this reason, we excluded data set 1 from our analyses.

#### Criteria for including neurons in analyses

For our analyses, we wanted to use cells that were visually responsive and well-tuned. For each data set, we applied the following to criteria to decide whether cells were included for analysis or not: Cells were visually responsive if they displayed an evoked response that was at least one standard deviation above or below their spontaneous rate. The average and standard deviation of the spontaneous rate were computed from the final 500 ms of the blank screen presentation in between trials. Cells were well-tuned if they were approximated by the sum of two Von mises functions with an *r*^2^ *>* 0.75 (Graf et al., 2011).

#### Measuring amplitude, bandwidth, and orientation tuning curves

For each cell, we first found the maximum value of the tuning curve. We then considered the half of the tuning curve where the maximum value was (0-175 deg or 180-355 deg), and convolved this with [0.05, 0.25, 0.4, 0.25, 0.05] for smoothing. We measured amplitude as the difference between the maximum and minimum values of the smoothed tuning curve. To measure bandwidth, we first found the two points on either side of the peak that were closest 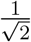 to of the peak value after baseline subtracting. Then, we defined the bandwidth as half of the difference in deg of two points. Because our analyses focused on *orientation* decoding, not direction of motion decoding, we used only the responses from the half of the tuning curve that contained the peak.

#### Upsampled pseudo-populations

For the neuronal data analysis, we created upsampled pseudo-populations as follows: For one orientation, say 0 degrees, we selected 5 neurons from the original population at random. Then, we shifted the 5 selected neurons’ responses to the stimuli such that their peak responses were at 0 degrees. We repeated this for each orientation from 0, 5, …, 175 degrees, creating a total population of 180 neurons.

#### Fisher information

Here we use the *linear* Fisher information, which assumes linear decoding and ignores the information available to nonlinear decoders. It is defined as follows for each stimulus, *θ* (Kafashan et al., 2021; Seriès et al., 2004).

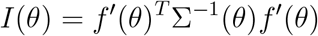

where *f* ^′^(*θ*) is a vector of the difference in the cells’ responses to *θ* and *θ* + 5 degrees, because the neuronal data was sampled at 5 degree spacing. Σ(*θ*) is the population’s covariance matrix across the trials for *θ*. We average *I*(*θ*) across all 36 orientations to get one value for the data set. The linear Fisher information is preferable to use for neuronal data because the total Fisher information depends on Σ^′^(*θ*), which is unstable without a large number of trials.

## Supporting Information

### Proof for the centroid norm increasing with amplitude heterogeneity

First consider the centroid norm of the manifold created by the responses to stimulus *θ* for a homogeneous population. The neurons have identical tuning curve shapes given by *f* (*θ*). The neurons have different peak preferences of stimuli, so the *i*th neuron’s tuning curve is given by *f*_*i*_(*θ*) = *f* (*θ* − *θ*_*i*_) where the *i*th neuron’s preferred stimulus is *θ*_*i*_. The firing rate for the *i*th neuron for stimulus *θ* is *m*_*i*_(*θ*) = *f*_*i*_(*θ*) + *s* where *s* is a fixed spontaneous firing rate held constant for all neurons.

The global mean response of each neuron is given by the average of its responses to all stimuli, 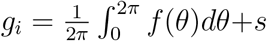. The centroid vector at stimulus *θ*_1_ is *C* = (*m* (*θ*), …, *m* (*θ*)). The the norm of the global mean subtracted centroid is given by:

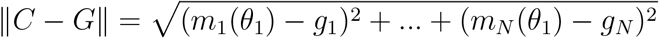

Now consider a population with heterogeneous amplitudes. Suppose for simplicity that heterogeneity is implemented by drawing *N* independent and identically distributed *ϵ*_*i*_ ∼ N(0, *κ*) (where *κ* indicates the level of heterogeneity), and scaling the *i*-th tuning curve by 1 + *ϵ*_*i*_. The heterogeneous tuning curves are given by 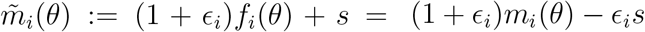.

The new global means are given by 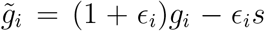 and the new centroid vector for the manifold for stimulus *θ*_1_ is given by 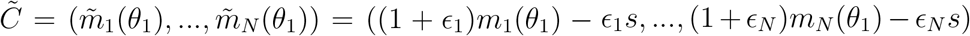 We can write the norm of the global mean subtracted centroid in this case as:

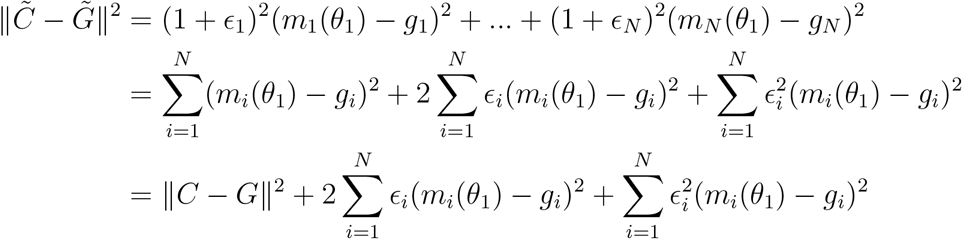

Because 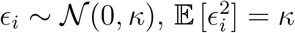. Hence,

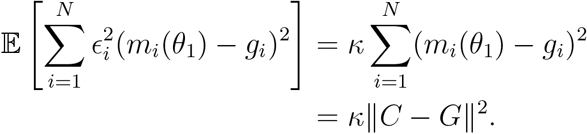

Additionally, because E [*ϵ*_*i*_] = 0,

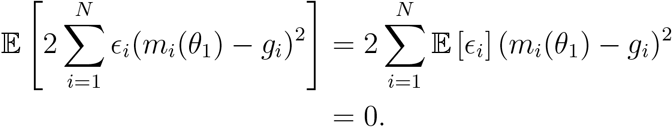

Taken together, the above equations show that on average 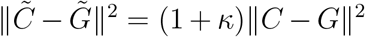, and the centroid norm increases with the heterogeneity level, *κ*.

**SI Table 1.** Multiple regression results for subpopulation properties as predictors of capacity. From data set 2, we created 500 subpopulations and 500 pseudo-populations. We measured the mean amplitude, standard deviation of amplitudes, mean bandwidth, and standard deviation of bandwidths for each of the subpopulations, and their capacity for discrimination and identification. For each task and type of subpopulation, we show the standardized regression coefficients for the mean and standard deviation of the tuning properties as a predictors of capacity.

| Task and population type | Amplitude |  | Bandwidth |  |
| --- | --- | --- | --- | --- |
|  | Mean | Standard deviation | Mean | Standard deviation |
| Discrimination, true | 0.30 | 0.080 | -0.74 | -0.13 |
| Discrimination, pseudo | 0.29 | 0.11 | -0.12 | -0.10 |
| Identification, true | 0.62 | -0.011 | -0.031 | -0.18 |
| Identification, pseudo | 0.34 | 0.31 | -0.067 | -0.16 |

**SI Table 2.**
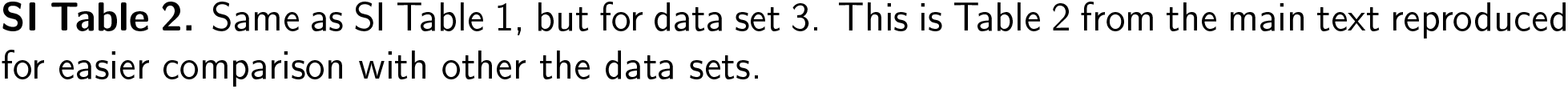
Same as SI Table 1, but for data set 3. This is Table 2 from the main text reproduced for easier comparison with other the data sets.

**SI Table 3.** Same as SI Table 1, but for data set 4.

| Task and population type | Amplitude |  | Bandwidth |  |
| --- | --- | --- | --- | --- |
|  | Mean | Standard deviation | Mean | Standard deviation |
| Discrimination, true | 0.44 | 0.14 | -0.37 | 0.09 |
| Discrimination, pseudo | 0.27 | 0.42 | -0.48 | 0.12 |
| Identification, true | 0.68 | -0.24 | -0.22 | -0.030 |
| Identification, pseudo | 0.34 | 0.40 | -0.40 | 0.60 |

**SI Table 4.** Same as SI Table 1, but for data set 5.

| Task and population type | Amplitude |  | Bandwidth |  |
| --- | --- | --- | --- | --- |
|  | Mean | Standard deviation | Mean | Standard deviation |
| Discrimination, true | 0.093 | 0.65 | -0.067 | -0.062 |
| Discrimination, pseudo | 0.022 | 0.59 | -0.14 | -0.18 |
| Identification, true | 0.50 | 0.24 | 0.15 | -0.025 |
| Identification, pseudo | 0.14 | 0.66 | -0.17 | -0.10 |

**SI Figure 1.**
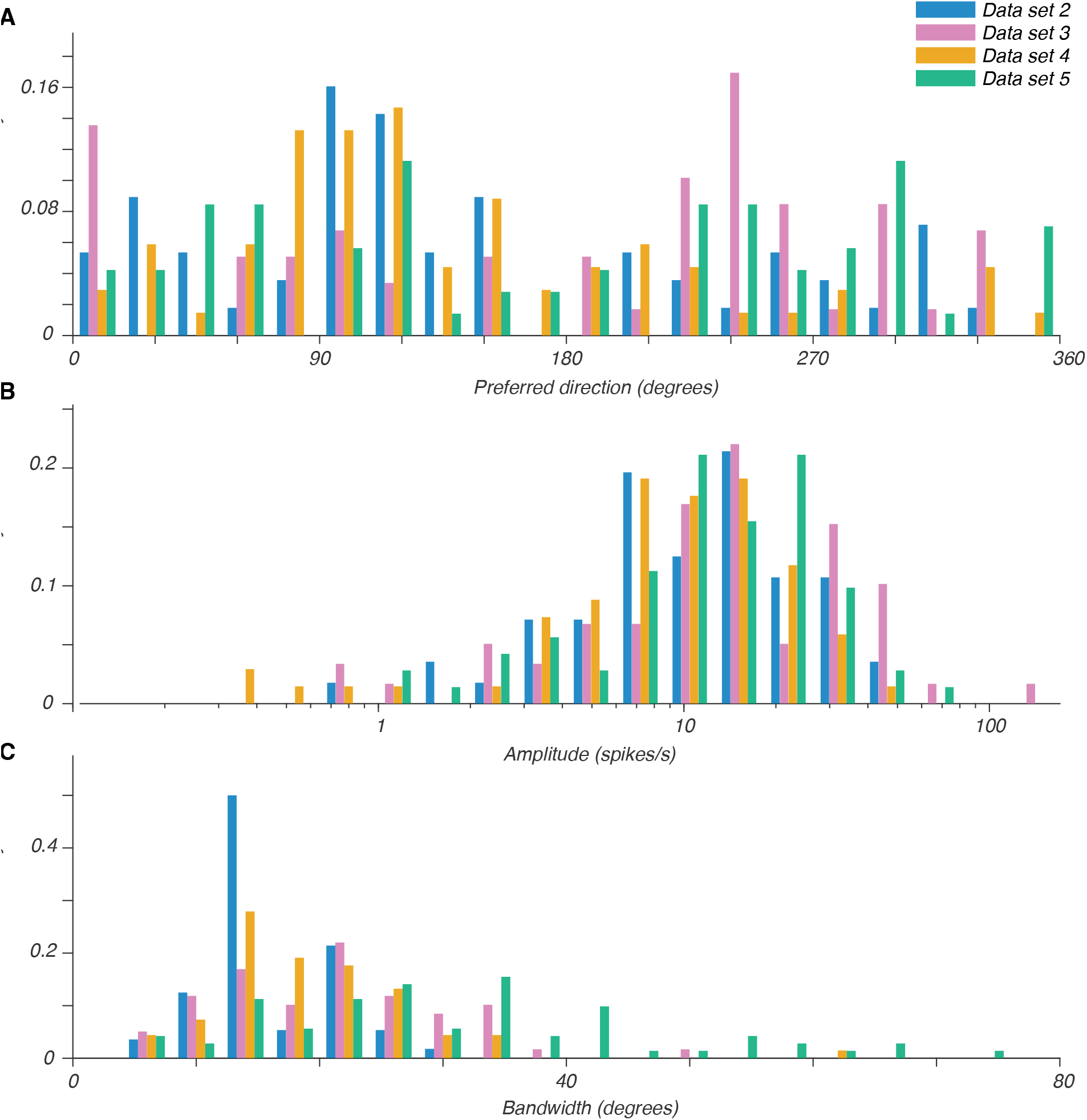
Histograms of **A** preferred directions, **B** amplitudes, and **C** bandwidths for all four datasets. Data set 2 had 56 neurons, data set 3 had 59, data set 4 had 68, and data set 5 had 71.

**SI Figure 2.**
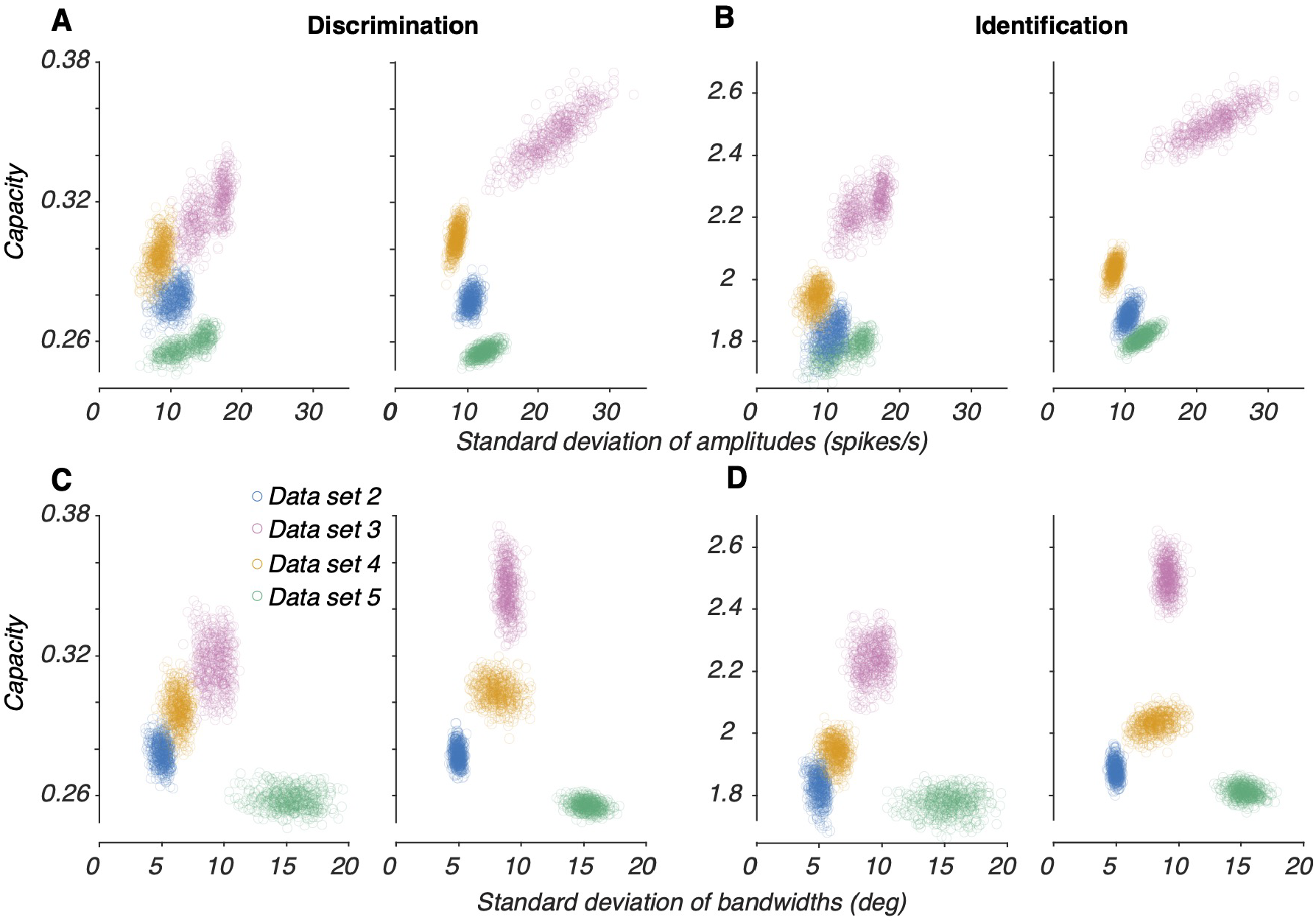
Relationship between diversity and capacity for all four data sets. Data set 3 is reproduced from Figure 4 for comparison with the other datasets. For each data set, we selected 500 subpopulations of 30 neurons each and 500 pseudo-populations of 180 neurons each. **A** Amplitude diversity versus capacity for discrimination in the true subpopulations (left) and pseudo-populations (right). **B** Same as panel A, but for identification. **C** Bandwidth diversity versus capacity for discrimination in the true subpopulations (left) and pseudo-populations (right). **D** Same as panel C but for identification.

